# Bed Nucleus of the Stria Terminalis GABA neurons are necessary for changes in foraging behavior following an innate threat

**DOI:** 10.1101/2023.02.25.530051

**Authors:** Annie Ly, Alexandra Barker, Emily D. Prévost, Dillon J. McGovern, Zachary Kilpatrick, David H. Root

## Abstract

Foraging is a universal behavior that has co-evolved with predation pressure. We investigated the role of bed nucleus of the stria terminalis (BNST) GABA neurons in robotic and live predator threat processing and their consequences in post-threat encounter foraging. Mice were trained to procure food in a laboratory-based foraging apparatus in which food pellets were placed at discrete and incrementally greater distances from a nest zone. After mice learned to forage, they were exposed to either a robotic or live predator threat, while BNST GABA neurons were chemogenetically inhibited. Post-robotic threat encounter, mice spent more time in the nest zone, but other foraging parameters were unchanged compared to pre-encounter behavior. Inhibition of BNST GABA neurons had no effect on foraging behavior post-robotic threat encounter. Following live predator exposure, control mice spent significantly more time in the nest zone, increased their latency to successfully forage, and their overall foraging performance was significantly a ltered. I nhibition o f BNST GABA neurons during live predator exposure prevented changes in foraging behavior from developing after live predator threat. BNST GABA neuron inhibition did not alter foraging behavior during robotic or live predator threat. We conclude that while both robotic and live predator encounter effectively intrude on foraging behavior, the perceived risk and behavioral consequence of the threats are distinguishable. Additionally, BNST GABA neurons may play a role in the integration of prior innate predator threat experience that results in hypervigilance during post-encounter foraging behavior.

## Introduction

From nematode to mammal and all remaining subjects in kingdom Animalia, foraging for food is a shared behavior [1-4]. Foraging for food encompasses a multitude of neural systems: hunger, satiety, sensorimotor integration, attention, navigation, exploration, and risk assessment to name a few [3]. While there are many factors involved in foraging behavior, a hunger drive exists to motivate food consumption that leads to satiation [5]. However, foraging is not a behavior that exists in a vacuum in which animals navigate environments to satiate hunger [6]. A limitation for most animals is that they are at risk of predation, and predator threat influences foraging behavior [7].

Predation provides a major evolutionary pressure for animals to develop highly conserved defensive behaviors while foraging. One example of a defensive foraging behavior seen across fish, birds, and mammals is an avoidance of open spaces [8]. For example, when a wooden model of a hawk was flown over black-capped chickadees, they preferred to carry their food back to shelter, rather than consume their food in the open, where there was a perceived predator threat [9]. Another instance of a defensive foraging behavior, particular to nocturnal animals, is the avoidance of bright light, such as natural moonlight and artificial light [10-14]. Avoidance is an especially crucial defensive behavior during foraging when the threat of predation is uncertain or ambiguous, particularly after a prior predator experience. Therefore, animals must evaluate between food attainment and predation risk [7,15,16].

The bed nucleus of the stria terminalis (BNST) is an extended amygdala structure that is strongly implicated in the processing of uncertain or ambiguous threats [17-22] and feeding [23-25]. We hypothesized that the BNST plays an important role in foraging behavior post-threat encounter when the threat is absent yet uncertain. The BNST is heterogeneous in neuronal cell-types but is a predominantly GABAergic brain region [26,27].

In order to investigate the role of GABA BNST neurons in foraging behavior after a threat, we used a chemogenetic approach to inhibit GABA BNST neurons during predator exposure and assessed behavioral changes in a foraging task following the predator experience. Many laboratory-based foraging paradigms exist for rodents [16,28-33]. Here, we compared the effects of a semi-natural “robogator” predator threat [31] and a live predator threat on post-encounter foraging behavior, as well as identified the role of GABA BNST neurons.

We found that both robogator and live predator threat disrupted post-encounter foraging behavior, but live predator threat was more efficacious. While chemogenetic inhibition of BNST GABA neurons during robogator threat had no effect on post-encounter foraging, inhibition of BNST GABA neurons during live predator threat prevented changes in foraging that were observed post-encounter. BNST GABA neuron inhibition did not affect foraging behavior during robotic or live predator threat. We have interpreted these results to show that while both robotic and live predator threat effectively intruded ongoing foraging behavior, live predator threat resulted in pronounced disruptions of post-encounter foraging more so than robotic threat. Additionally, our results have indicated that BNST GABA neurons may play a role in the integration of prior innate predator threat experience with hypervigilance as a behavioral consequence.

## RESULTS

### Baseline foraging behavior in a novel environment

Throughout the foraging sessions, mice were trained to retreat back to the nesting zone after consuming a food pellet at discrete and incrementally greater distances from the nesting zone (**Figure 1A**). In the first several sessions, mice did not successfully forage equally across the distances of the food pellet location. Compared to the 30cm trial, mice took significantly longer to successfully forage at the 50cm [hazard ratio (HR) = 0.63; 95% confidence interval (CI) = 0.53-0.77; p<0.001] and 70cm [HR = 0.65; 95% CI = 0.54-0.79; p<0.001] distances (**Figure 1B**). For the first 12 consecutive sessions, food consumption latency varied with a session-sex interaction effect [F(11,326) = 26.9, p = 0.005]. In the last 4 consecutive sessions, there was no significant difference in behavior due to sex, nor a significant interaction with session or trial (**Figure 1C)**. A main effect of trial remained on foraging latency, but this effect is likely attributed to the increasing distances the mice must traverse away from the nesting zone to obtain the food pellet [main effect (trial): F(2,185) = 16.9, p = 0.0002]. There was a significant difference across the first 12 consecutive foraging sessions for total time spent in the foraging zone [main effect (session): F(11,338) = 85.4, p<0.0001]. This effect is likely due to the mice spending increasingly more time in the foraging zone rather than the nesting zone before procuring and consuming the food pellet. In the last 4 consecutive sessions, no significant sex difference nor a significant interaction with session or trial was observed with time spent in the foraging zone (**Figure 1D)**. Lastly, there was a significant difference in number of nest exits across the first 12 consecutive foraging sessions [main effect (session): F(11,338) = 72.9, p<0.0001]. As mice learned to forage over days, the number of nest exits were significantly reduced. In the last 4 consecutive sessions, there was no significant sex difference nor an interaction with session or trial, regarding number of nest exits (**Figure 1E)**. Thus, by the end of training, male and female mice learned to traverse the foraging zone to retrieve the food pellets.

**Figure 1.**
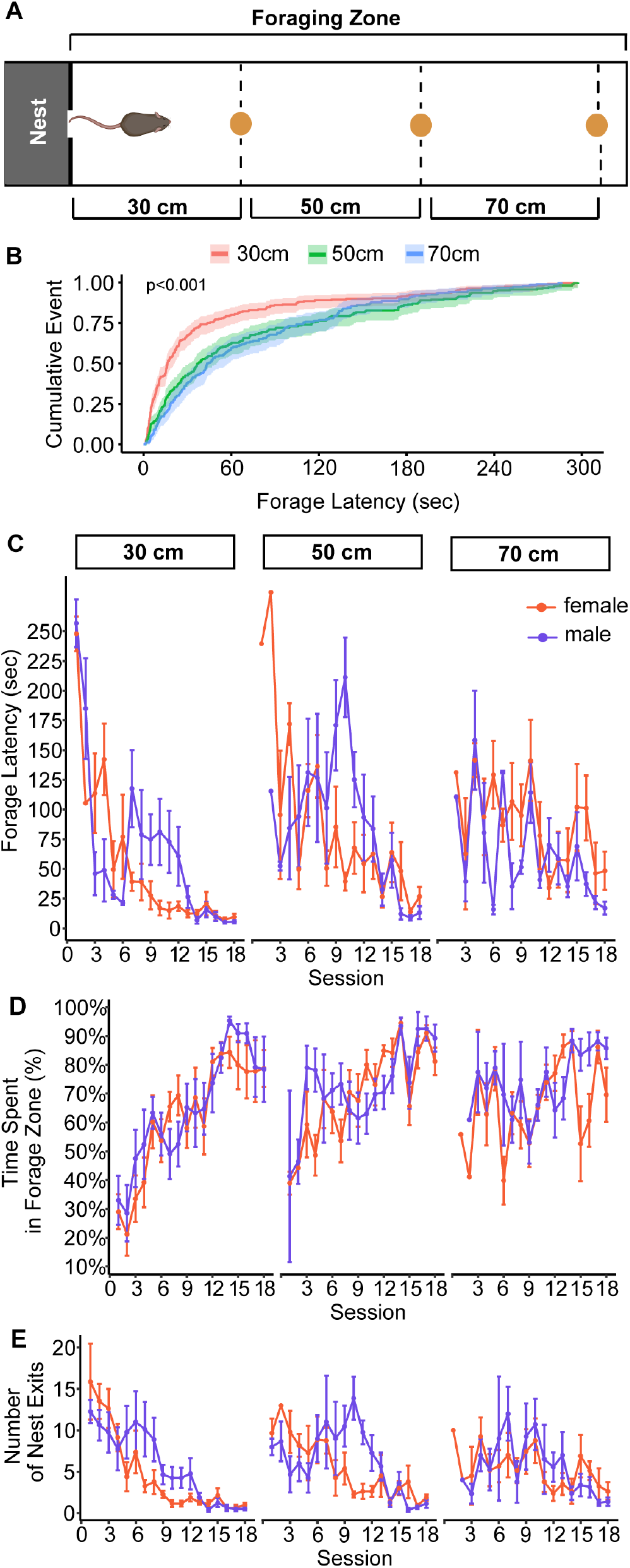
Foraging performance of mice in a novel environment. (A) Illustration of the foraging apparatus and placement of the food pellet for each trial in a session. (B) Significant differences in latency to successfully forage by trial, illustrated by Cox-proportional hazards regression model. No significant sex differences in the last 4 consecutive sessions were observed for (C) foraging latency, (D) time spent in foraging zone, and (E) number of nest exits.

### Robotic and live predator threats produce similar changes in foraging behavior, but BNST GABA inhibition during imminent predator threat exposure has no effect on foraging

Prior to testing the role of BNST GABA neurons in the interaction of threat and foraging behavior, we tested whether hM4D(Gi) was capable of reducing BNST GABA neuronal activity. VGaT-IRES::Cre mice were bilaterally injected in BNST with Cre-dependent adeno-associated viruses encoding the inhibitory designer receptor (hM4D(Gi)) or fluorophore control (GFP) to exclusively transfect BNST GABA neurons (**Figure 2A-C**). GFP and hM4D(Gi) mice were then intraperitoneally injected with a behaviorally-subthreshold dose of clozapine (0.1 mg/kg) [41] to chemogenetically inactivate BNST GABA neurons at least 10 minutes prior to receiving five footshocks (**Figure 2D**). Five footshocks were selected as the method for chemogenetic inhibition validation based on prior work showing that aversive stimuli reliably activate BNST neurons [42,43]. Mice were euthanized and perfused 90 minutes later, and virally-labeled BNST neurons were evaluated for c-Fos co-expression. BNST GABA neurons with hM4D(Gi) co-expressed significantly less c-Fos than GFP BNST GABA neurons [t(6) = -7.45, p = 0.005] (**Figure 2E**). Thus, our results indicate that hM4D(Gi) can reduce BNST GABA activity.

**Figure 2.**
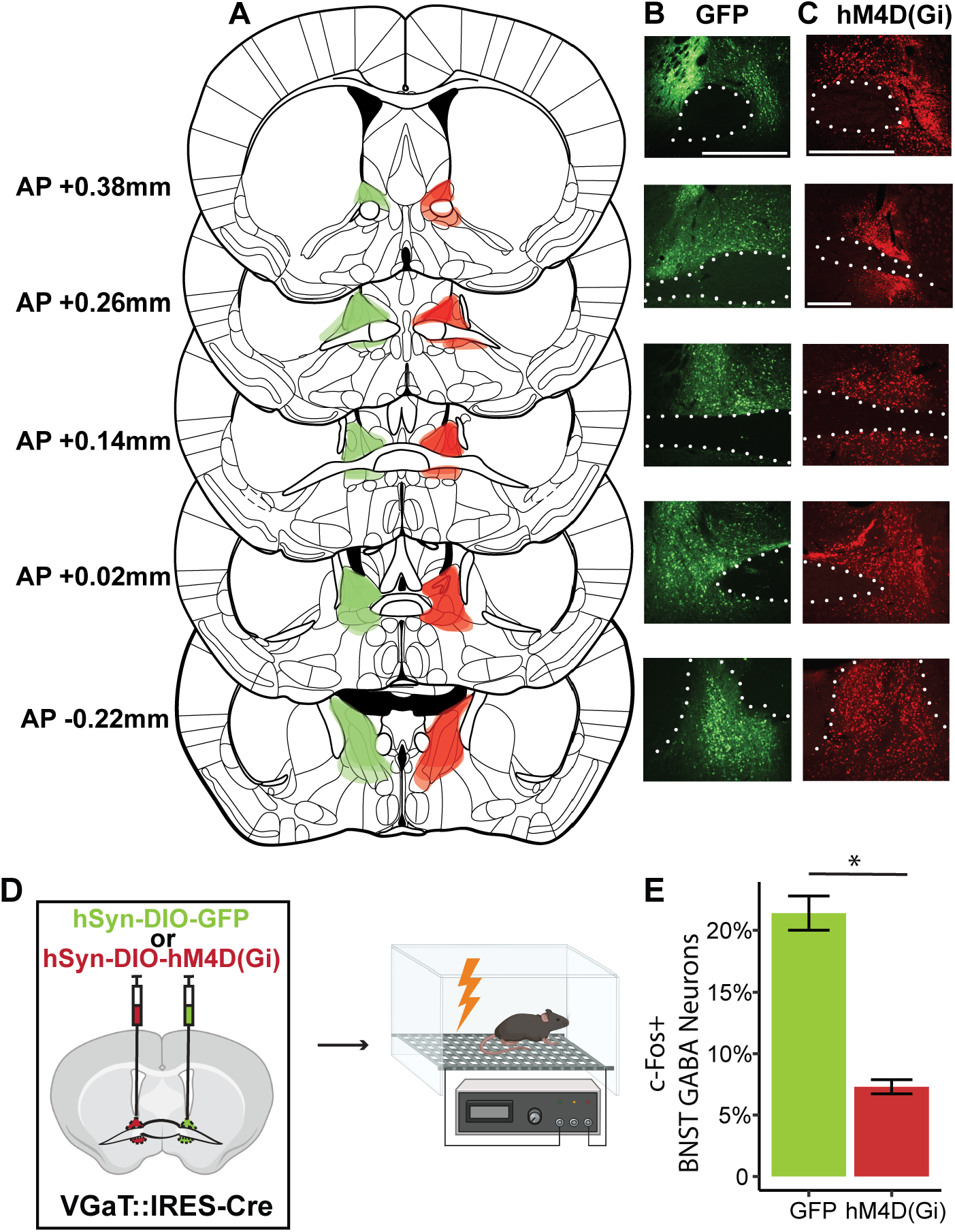
Histological verification of chemogenetic expression and targeting. (A) Schematic depicting localization of fluorescence across the BNST. Representative images of each anteroposterior (AP) coordinate for (B) GFP in green and (C) hM4D(Gi)-mCherry in red. Scale bar=100μm. (D) Experimental depiction of chemogenetic verification by 5 0.5mA footshocks. (E) Significantly fewer hM4D(Gi)-labeled BNST GABA neurons co-expressed with c-Fos than GFP-labeled BNST GABA neurons.

After baseline foraging behavior was achieved, mice were injected with clozapine (0.1mg/kg) to chemogenetically inactivate BNST GABA neuron signaling and challenged to forage under robotic or live predator threat (**Figure 3A**). Foraging behavior under a predatory threat changed on multiple parameters. The latency to successfully forage by consuming the food pellet was also significantly increased under both predator threats [main effect (pre- and circa-encounter): F(1,56) = 4.51, p = 0.03] (**Figure 3B**). Additionally, mice spent significantly more time in the nest circa-predator exposure [main effect (pre- and circa-encounter): F(1,56) = 8.51, p = 0.005], regardless of the predatory threat being live or robotic (**Figure 3C-D**). Similarly, mice made more exits from the nest during predator threat, but there was no difference between live or robotic threat [main effect (pre- and circa-encounter): F(1,56) = 9.36, p = 0.003] (**Figure 3E-F**). While robotic and live predator threats similarly changed foraging behavior, chemogenetic inactivation of BNST GABA neuron signaling failed to change any foraging behavior during the threat when the predator was present.

**Figure 3.**
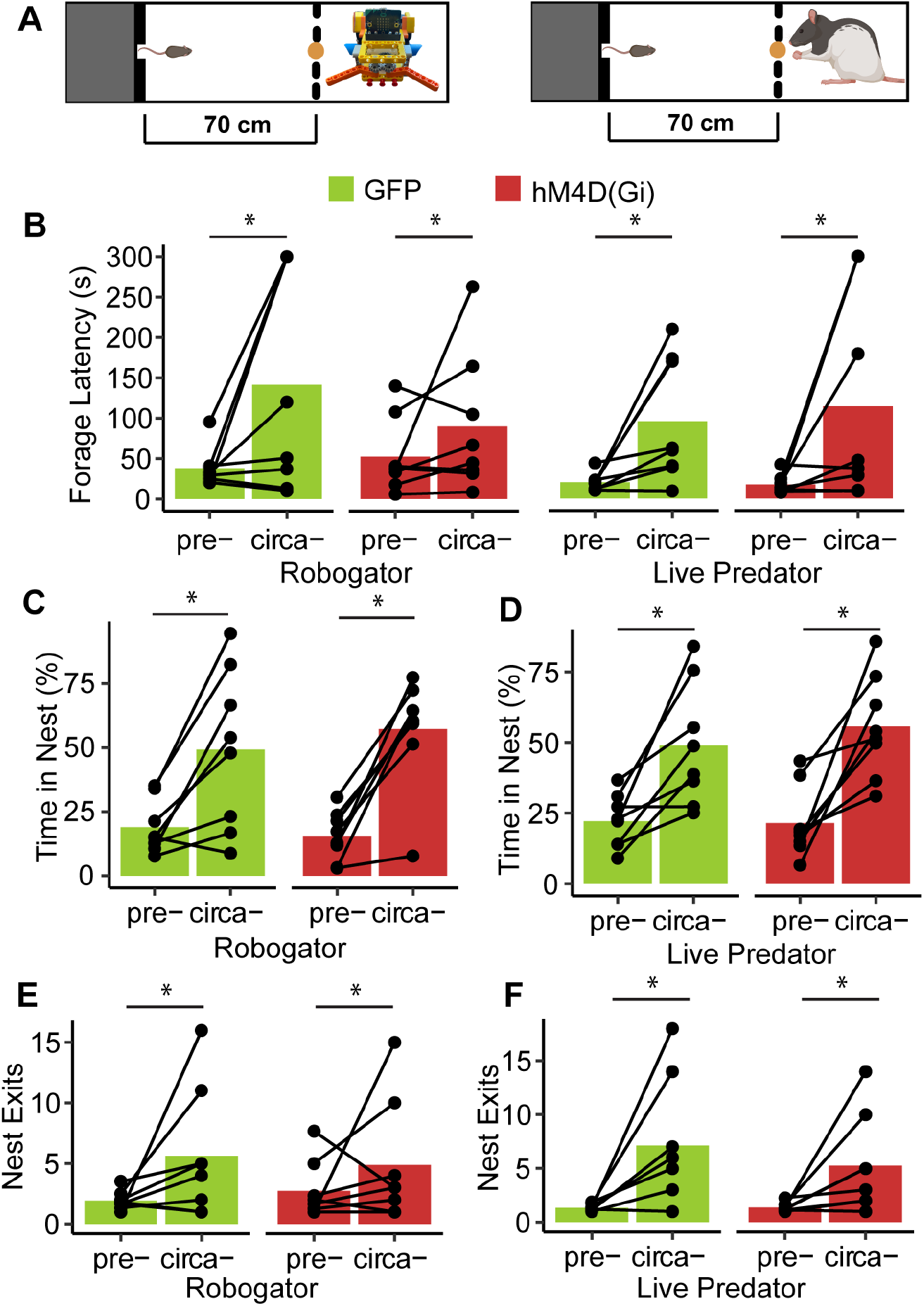
Foraging behavior in the presence of a predator threat with chemogenetic inactivation of BNST GABA neuron signaling. (A) GFP and hM4D(Gi) mice were injected with clozapine and exposed to a predator threat. (B) Foraging latency increased circa- robogator and live predator. Time spent in nest increased circa- (C) robogator and live predator. Nest exits increased circa- (E) robogator and (F) live predator. (G) The first latency to exit the nest increased circa- robogator and live predator, and mice took longer to exit the nest with a live predator present than a robotic predator.

### BNST GABA inhibition does not change foraging behavior following robotic predator exposure

After predator exposure and chemogenetic inhibition, we examined foraging behavior 48-hours post-encounter in the absence of threat (**Figure 4A**). For the robotic predator, across all trials, there was no significant effect of predator encounter or chemogenetic inactivation of BNST GABA neurons on foraging latency (**Figure 4B**). However, mice spent significantly more time in the nest after robotic predator regardless of BNST GABA inactivation [main effect (encounter): F(1,236) = 15.4, p<0.001] (**Figure 4C**). We next examined multiple foraging parameters, such as foraging latency, number of nest exits, time spent in nest, distance traveled, first nest exit latency, and last nest exit latency. Each variable was normalized on a scale of 0 to 1 by robogator encounter (pre-, post-) and by trial (30cm, 50cm, 70cm) for GFP and hM4D(Gi) mice (**Figure 4D-E**). Across all foraging parameters by calculating the polygon area of each mouse before and after robotic predator, there was no significant effect of either robogator or chemogenetic inactivation of BNST GABA neurons (**Figure 4F**). These results suggest that time spent in nest after an encounter with the robotic predator was the primary behavior change observed. Additionally, any post-encounter foraging changes were not augmented with BNST GABA inhibition.

**Figure 4.**
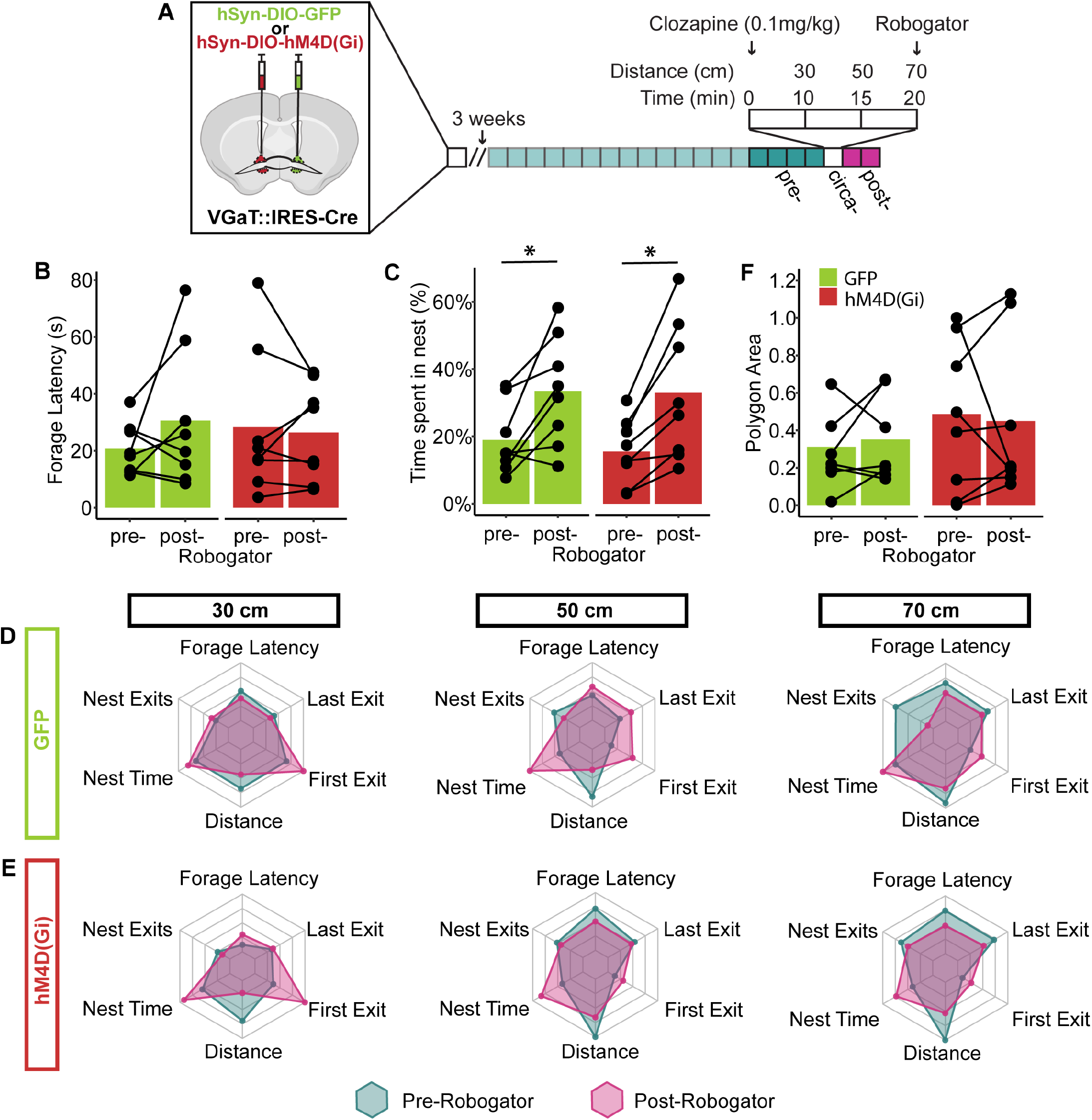
Foraging performance post-robogator encounter and chemogenetic inactivation of BNST GABA neuron signaling. (A) Experimental timeline. (B) No significant differences in foraging latency by treatment or robogator encounter. (C) Mice spend more time in the nest after robogator encounter, regardless of chemogenetic inactivation. Foraging performance by trial before and after robogator exposure for (D) GFP mice and (E) hM4D(Gi) mice. The maximum axis for each radar plot is equal to the maximum average out of the foraging parameters. (F) No changes in overall foraging performance by treatment or robogator encounter.

### BNST GABA inhibition prevents post-live predator changes in foraging

After assessing robogator post-encounter foraging behavior, mice were re-trained to forage with no group differences in foraging latency before being exposed to a live predator (**Figure 5A-B)**.In contrast to robogator threat, foraging latency following live predator threat depended on the interaction between live predator exposure and BNST GABA neuron inactivation [main effect (encounter): F(1,334) = 16.9, p<0.001; encounter x treatment, F(1,332) = 5.93, p = 0.01] (**Figure 5C**). In the absence of a live predator, but after the experience of encountering one, GFP mice showed significantly increased foraging latency, and this effect was absent in hM4D(Gi) mice [comparisons of estimated marginal means of pre-x GFP versus post-x GFP, p = 0.0003]. Time spent in nest also showed a significant interaction between live predator exposure and BNST GABA neuron inactivation [main effect (encounter): F(1,334) = 7.04, p = 0.008; encounter x treatment, F(1,332) = 12.9, p<0.001] (**Figure 5D**). GFP mice showed significantly increase time in nest post-encounter, and this effect was blocked in hM4D(Gi) mice [comparisons of estimated marginal means of pre-x GFP versus post-x GFP, p = 0.04].

**Figure 5.**
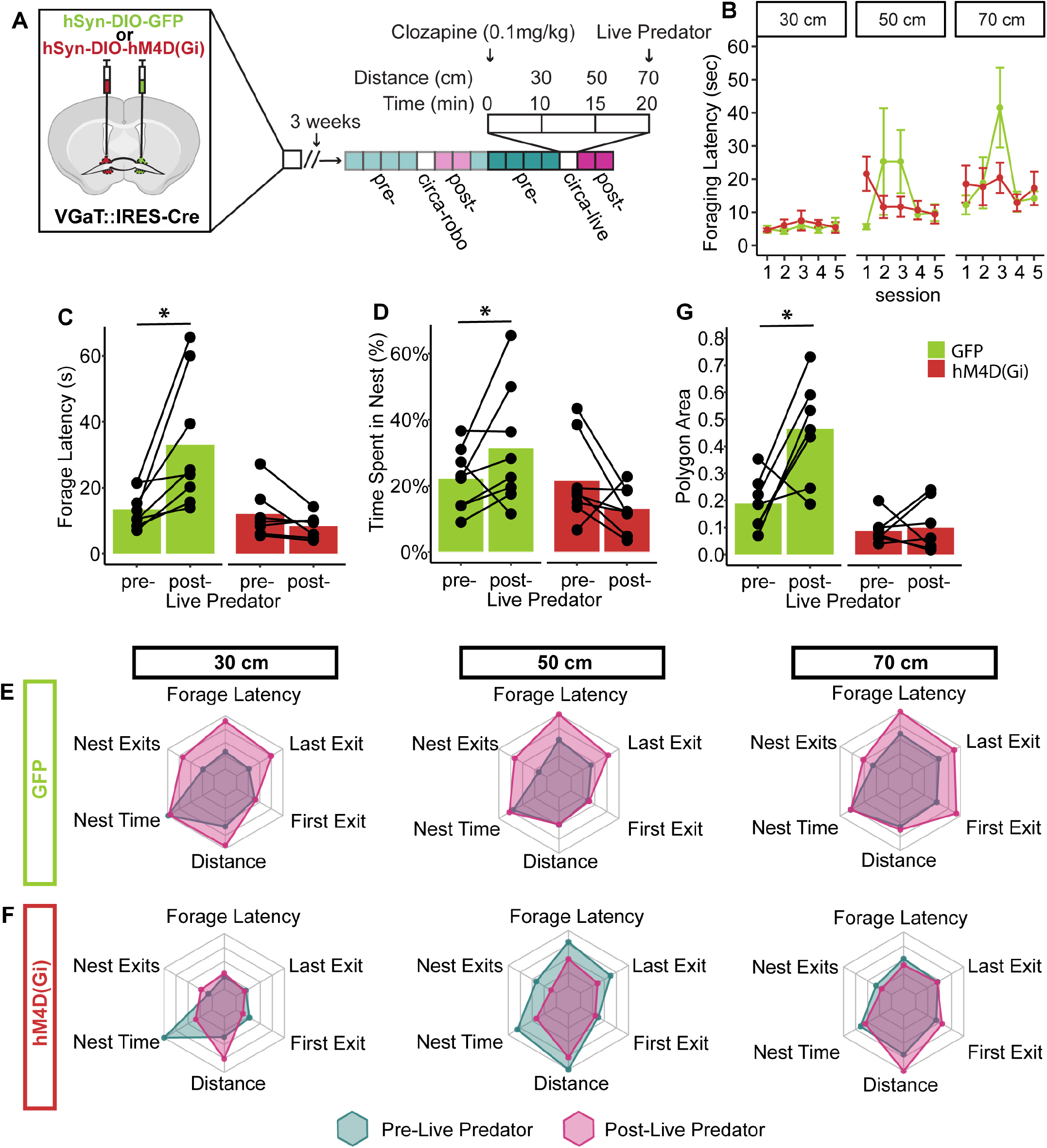
Foraging performance post-live predator encounter and chemogenetic inactivation of BNST GABA neuron signaling. (A) Experimental timeline. (B) No treatment group differences in foraging latency pre-live predator encounter. Significant effect of chemogenetic BNST GABA inactivation on (C) foraging latency and (D) time spent in nest. Foraging performance by trial before and after live predator exposure for (E) GFP mice and (F) hM4D(Gi) mice. (G) Significant effect of chemogenetic BNST GABA inactivation on overall foraging performance after live predator exposure.

To understand the effect of predator exposure and BNST GABA chemogenetic inactivation on multiple foraging parameters, we normalized selected foraging parameters (foraging latency, last nest exit, first nest exit, distance traveled, nest time, nest exits) on a scale of 0 to 1 within foraging parameter and trial (30cm, 50cm, 70cm). We defined overall foraging performance as the polygon area calculated from the shape of all parameters plotted on a polar plot. Across all foraging parameters, we found that live predator exposure had a significant effect on overall foraging performance post-encounter that depended on an interaction between live predator exposure and chemogenetic inactivation of BNST GABA neurons [main effect (encounter): F(1,458) = 551.5, p<0.001; encounter x treatment, F(1,456) = 238.5, p<0.001] (**Figure 5E-G**). Post-hoc analysis identified that the foraging performance of GFP mice was significantly different between the pre- and post-live predator encounter, and this effect was not observed in hM4D(Gi) mice [comparisons of estimated marginal means of pre-x GFP versus post-x GFP, p<0.0001]. These results of foraging behavior 48-hours after the live predator encounter suggest that BNST GABA inactivation prevented significant changes that are normally the result of prior threat experience.

### Locomotor path does not change after predator exposure and BNST GABA inhibition

Confrontation with a predator threat can change the distance traveled and trajectory path of the prey [31,44,45]. Here, we assessed changes in locomotor activity to determine the acute effects of predator exposure and BNST GABA inhibition on foraging. Based on linear modeling of the tracked center point of the mice, there was no effect of predator exposure on the estimated linear path taken while foraging [r(118) = 14.3, p = 0.57]. Neither was there a difference between trials [r(118) = 5.24, p = 0.77]. Overall, there was also no interaction effect of predator exposure and BNST GABA inhibition on locomotor path [r(118) = -10.6, p = 0.50] (**Figure 6A-D**). Similarly to robotic predator exposure, there was no effect of live predator exposure [r(108) = 16.4, p = 0.54], trial [r(108) = 8.67, p = 0.66], or interaction effect of live predator and BNST GABA inhibition [r(108) = -11.7, p = 0.50] on locomotion afterwards (**Figure 6E-H**). In summary, chemogenetic inhibition of BNST GABA neurons did not alter post-encounter locomotor path for neither robotic nor live predator experience.

**Figure 6.**
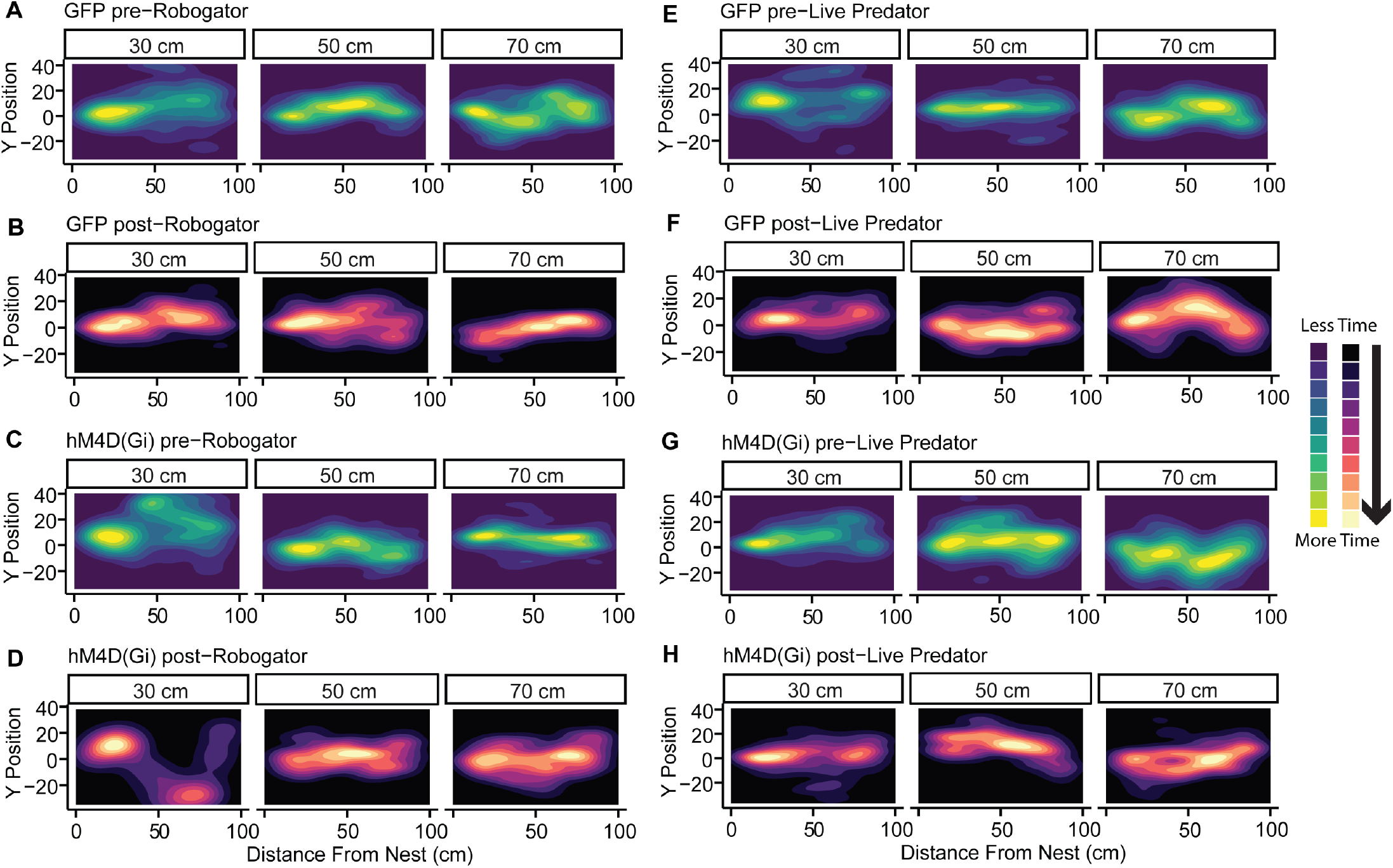
Locomotor path patterns before and after predator threat exposure. Averaged linear path taken by GFP mice (A) before robogator and (B) after robogator. Averaged linear path taken by hM4D(Gi) mice (C) before robogator and (D) after robogator. Averaged linear path taken by GFP mice (E) before live predator and (F) after live predator. Averaged linear path taken by hM4D(Gi) mice (G) before live predator and (H) after live predator.

## DISCUSSION

Predation risk not only impacts foraging behavior in the presence of a predator but also in the immediate aftermath of a close encounter. Using a laboratory-based foraging task, we evaluated foraging performance before, during, and after two distinct predator threats, as well as identified the role of BNST GABA neurons in modulating these behaviors. During initial foraging training, mice took significantly longer to procure food at greater distances away from the nest. The increased foraging latency is consistent with an innate avoidance of bright, open spaces in rodents, concomitant with an increasing distance from a dark enclosure [8,10-14]. Importantly, by completion of training, mice were able to efficiently navigate and retrieve the food pellet.

Previous research in laboratory rats observed sex differences in foraging strategy involving male rats spending more time in open, novel environments compared to female rats [32,44,46,47]. We found no sex differences in mice across foraging success, foraging latency, time spent in the foraging zone, and nest exits. While there was a session-sex interaction effect in the first 12 sessions of initial training on foraging latency, there were no interactions or main effects of sex following extensive training.

It is well established in field studies that animals change their foraging behavior under predation risk [7,8,15]. Particularly, they will flee when a predator reaches a distance that is perceived as threatening to survival, a concept coined by ethologists as the flight-initiation distance [48,49]. Fanselow and Lester articulate the psychological perception of threat in their predator imminence continuum theory [50]. One advantage of robotic predator threat is the ability to program a reliable predator-prey interaction, thus making the stimulus consistent across subjects [31]. Here, we found that foraging behavior was significantly disrupted during robotic and live predator threat. Others have revealed similar findings using either a robotic or live predator threat to show interference in successful foraging [33,34,51]. Importantly, we found no difference in time spent in nest, foraging latency, and nest exits between robogator and live predator threats, suggesting that a robotic predator threat is just as effective as a live predator in disrupting foraging behavior. However, chemogenetic inactivation of BNST GABA neuron signaling had no effect on any foraging behavior during robotic and live predator threat. Thus, BNST GABA neuron signaling does not appear to be involved in the circa-strike threat response to either a robotic or live predator.

Predator-prey interaction models suggest that even a brief predator exposure after chronic low-risk foraging may result in an overestimated defensive behavior response, such as hypervigilance [52]. In fact, a brief predator exposure can be followed by a long latency period before animals resume activity levels comparable to before the predator encounter [53]. Post-encounter, we found that there was no significant change in foraging latency with the robotic predator threat, but mice spent significantly more time in the nest. Others have reported similar observations after a robotic predator experience; while not exactly measuring time spent in nest, rats appeared to display far more “pause-then-retreat” behavior rather than “pause-then-approach” after a robotic predator encounter [54]. Pauses are likely part of a repertoire of defensive behavior to better assess uncertain risk of predation [33,55]. Spending more time in the nest while foraging latency remained unchanged suggests that the mice were spending more time in a safe zone to avoid predation risk. However, while robotic predator threat significantly increased time spent in nest, it did not impact overall foraging performance. It is possible that other measures, such as food intake over time, could be affected as others have shown [56]. Chemogenetic inactivation of BNST GABA neurons had no effect on the increased time spent in the nest post-robotic threat encounter.

In contrast to robotic predator, post-live predator encounter resulted in a constellation of changes in foraging behavior. Following live predator encounter, control mice showed prolonged foraging latency, increased time spent in the nest, and an overall reduction in foraging performance compared to pre-encounter levels. However, mice with BNST GABA neuron inhibition during live predator exposure showed no change in foraging latency, time spent in the nest, or overall foraging performance in post-encounter. Thus, inactivation of BNST GABA neurons was sufficient to rescue these behavioral changes.

When comparing between live and robotic predator threat, our results suggest that the robotic threat was effective in disrupting foraging behavior circa-encounter; however, it was not as salient of a threat experience as a live predator post-encounter. Why a robotic predator threat of this design may be perceived differently from a live predator may be due to a number of reasons; the robogator moved in a predictable manner or it lacked the innately aversive predator odor. This data indicate that robotic predator threat may not be sufficient to recruit BNST GABA neurons, and thus, BNST GABA neurons were not required for the increased time spent in the nest post-encounter. In contrast, with its great impact on multiple foraging parameters, live predator may present an innate threat than robotic predator. Indeed, rat aggression towards mice is observed as predatory behavior [57-59]. In addition, close proximity to a rat elicits a flight response in mice [60,61]. Thus, this innate threat experience required BNST GABA neurons for the consequential behavioral response in the post-encounter when the predator was no longer present.

Dynamic changes in locomotor path in a foraging task are dependent on GABA neuron signaling in the amygdala [31]. Specifically, regardless of overt threat, firing rates of single amygdala neurons are correlated with movement velocity in a foraging task [51]. We analyzed the locomotor path pre-encounter and post-encounter of robotic and live predator threat to examine the involvement of BNST GABA neurons in locomotor behaviors under threat. We found no effect of predator exposure or influence of BNST GABA neurons on locomotor path. The lack of a significant alteration in locomotor path as a result of predator experience might be explained by the extensive foraging training mice received in our paradigm, which was 18, compared to 2-7 foraging baseline sessions with rats [31,51]. Alternatively, the lack of BNST GABA neuron influence on locomotor path suggests that BNST GABA neuron signaling plays no role in foraging path after predator threat. This would support prior research demonstrating that extended amygdala regions, subdivisions, and cell-types may have specialized functions in behavioral output and threat processing [62-69]. In fact, the previous aforementioned experiments were of the lateral amygdala [31] and the basolateral amygdala [51], respectively.

In summary, we found that live predator threat induced more pronounced changes in post-encounter foraging than a robotic predator threat, and BNST GABA neurons were required in the post-live predator encounter changes in foraging. A parsimonious interpretation of the difference between robotic and live predator experience in eliciting changes in foraging behavior is that not all predator threat experiences are perceived as equal. Consistent with this interpretation, animals respond differently to predator species depending on the perceived psychological risk [48,49]. Given that GABA BNST activation is sufficient to produce anxiety-like behavior in mice [68,70,71], it is conceivable that BNST GABA neurons are required to induce an anxiety-like phenotype post-live predator threat encounter.

One limitation of the present study was that the robotic predator threat was followed by the live predator threat. While we cannot completely discount the influence of an order effect as an influence on BNST neurons, an order effect would have been more readily observed during the time of chemogenetic activation in the presence of a predator threat. However, chemogenetic inactivation of BNST GABA neurons had no effect on circa-threat foraging behavior for both robotic and live predator threats. In addition, following robotic threat, mice received further baseline foraging training that showed no foraging behavior differences between groups.

The observations made here may shed more light on the ethological behaviors of mice that distinguish the multiple components of the predator imminence continuum theory: pre-encounter, circa-strike, and post-encounter, as well as the role of the BNST, an extended amygdala structure, on discrete characteristics of a predator threat experience.

## MATERIALS AND METHODS

### Animals

VGaT-IRES::Cre knock-in mice (4-5 months old; 12 females,12males;B6J.129S6(FVB)-Slc32a1tm2(cre) Lowl/MwarJ; Stock #028862) were purchased from The Jackson Laboratory (Bar Harbor, ME) and bred at the University of Colorado Boulder (n=24). Mice were group-housed by sex (4-5 mice/cage) under a reversed 12hr:12hr light/dark cycle (lights on at 10pm) with access to water ad libitum. 16 mice were used for foraging behavior experiments, and 8 mice were used for c-Fos histology. For the duration of the foraging experiments, mice were weighed daily and fed to maintain 85% of their body weight. Food-restricted mice were fed after the foraging task. All experiments were performed during the dark phase of the light cycle. The experiments described were conducted in accordance with the regulations by the National Institutes of Health Guide for the Care and Use of Laboratory Animals and approved by the Institutional Animal Care and Use Committee at the University of Colorado Boulder.

### Surgery

Mice were anesthetized in a gasket-sealed induction chamber at 3% isoflurane gas. After confirming a surgical plane of anesthesia, isoflurane was continuously delivered at 1-2% concentration while the mouse was secured in the stereotactic instrument. AAV8-hSyn-DIO-hM4D(Gi)-mCherry (Addgene, n=8, 4 male, 4 female) or AAV8-hSyn-DIO-GFP (Addgene, n=8, 4 male, 4 female) were injected bilaterally into the BNST (5 × 1012 titer, 350 nL volume per hemisphere; 100 nl/min rate; +0.3 mm anteroposterior, ±0.6 mm mediolateral, -4.1 mm dorsoventral coordinates from bregma) using an UltraMicroPump, Nanofil syringes, and 35-gauge needles (Micro4; World Precision Instruments, Sarasota, FL). Syringes were left in place for 10 min following injections and slowly withdrawn. Mice were given 3 days of postoperative care and allowed 3-4 weeks of recovery before experimentation.

### Histology

Mice were anesthetized with isoflurane and perfused transcardially with 0.1M phosphate buffer followed by 4% (w/v) paraformaldehyde in 0.1M phosphate buffer, pH 7.3. Brains were extracted and cryoprotected in 18% sucrose solution in 0.1M phosphate buffer at 4°C overnight. Brains were cryosectioned to obtain coronal slices with BNST (30 μM). These coronal brain slices were mounted onto gelatin-coated slides and imaged for GFP or mCherry fluorescent expression on a Zeiss widefield Axioscope.

For c-Fos histology, mice were exposed to five footshocks (0.5mA, 500ms, 60s inter-shock interval) and perfused 90 minutes afterwards. As part of the perfusion protocol, mice were anesthetized with isoflurane and perfused transcardially with 0.1M phosphate buffer followed by 4% (w/v) paraformaldehyde in 0.1M phosphate buffer, pH 7.3. Brains were extracted and cryoprotected in 18% sucrose solution in 0.1M phosphate buffer at 4°C overnight. Brains were cryosectioned to obtain coronal slices with BNST (30 μM). Brain sections containing BNST were incubated with blocking buffer solution (4% bovine serum albumin and 0.3% Triton X-100 in 0.1M phosphate buffer (PB), pH 7.3) for 60 minutes, followed by incubation with mouse anti-GFP (1:500, Takara Bio, 632380), rabbit anti-mCherry (1:500, Takara Bio, 632496), and guinea pig anti-c-Fos (1:500, Synaptic Systems 226308) at 4°C overnight. Sections were washed in PB and incubated with donkey antimouse Alexa488 (1:200, Jackson ImmunoResearch, 715545150), donkey anti-rabbit Alexa594 (1:200, Jackson ImmunoResearch, 711585152), and donkey anti-guinea pig (1:200, Jackson ImmunoResearch, 706605148) for 120 minutes. These coronal brain slices were mounted onto gelatin-coated slides, coverslipped with ProLong DAPI diamond mounting medium (Invitrogen, P36971), and imaged for GFP, mCherry, and c-Fos fluorescent expression on a Nikon A1R confocal (20X). All GFP-positive, mCherry-positive, and c-Fos co-expressing cells within the BNST were counted in Adobe Photoshop between +0.38mm and -0.22mm from bregma. Cells were counted by scorers who were blinded to condition, and cells were only counted when it was also DAPI-positive.

### Foraging apparatus

A custom-made apparatus was built with a foraging zone (100cm length x 25cm width x 40cm height) and a nesting zone (15cm length x 25cm width x 40cm height), which were separated by a black Plexiglas gate that could be lifted up and down. The interior of the foraging zone was painted white, while the nesting zone remained as black Plexiglas. LED strip lights around the foraging perimeter maintained brightness at 40 lux, while the interior of the nesting zone was approximately 4 lux. A camera was positioned above the foraging apparatus to generate a video that was used with ANY-maze software (30hz, Stoelting Co. Wood Dale, IL) to track the subject’s center point in real time.

### Robotic and live predator

A four-wheel robot car (Shenzhen Yahboom Technology Co., Ltd.) with a programmable Micro:bit V1.5 board, a mechanical front-facing claw, and an infrared motion sensor module was assembled to mimic a previously described robotic predator design, herein referred to as robogator [31,34]. The dimensions are 24cm length x 15cm width x 12.5cm height. The Micro:bit board was programmed using the Microsoft MakeCode online platform. The robotic predator was programmed to detect a moving object at distance of less than 10cm. Once motion was detected, the LED screen of the Micro:bit board flashed 3 times before the robotic predator surged forward 20cm, snapped its mechanical claw 3 times, and retreated back to its original location.

A Long-Evans female rat (8 months old) was placed inside a clear plastic cage (28cm length x 15cm width x 23cm height) with a small circular opening on one side, and the filter on the cage top was removed. The rat was able to move freely within the cage to react to the presence of the mouse but was confined to prevent direct interaction.

### Behavioral Procedure

#### Foraging Baseline

Mice underwent 2 days of habituation to the nesting zone only of the foraging apparatus for 15 minutes with some home cage nesting material and 20 food pellets (45 mg, grain-based; F0165, Bio-Serv).

After habituation, mice were allowed to explore the foraging zone and consume a food pellet located at discrete, incremental distances. A single foraging session consisted of 3 trials, defined by the distance of the food pellet in relation to the nesting zone entrance: 30cm, 50cm, and 70cm. Every trial began with the opening of the Plexiglas gate and ended with the mouse back in the nesting zone after consuming the pellet. When a mouse consumed the pellet and returned back to the nesting zone, the Plexiglas gate was lowered, and the mouse was rewarded an additional food pellet in the nesting zone. The mouse was given 1 minute before the next trial began, and the Plexiglas gate was lifted. If a mouse failed to retrieve a food pellet within 5 minutes in any trial, the session would end with the mouse not receiving another opportunity to forage until the next session. Mice underwent training until there was no session effect on foraging latency across the last 4 consecutive sessions. Therefore, mice underwent 18 baseline foraging sessions.

#### Foraging circa- and post-<u>Robotic Predator Exposure</u>

Mice were exposed to the robotic predator in a single session in the last trial, in which the food pellet was located 70cm away from the nesting zone. Mice were intraperitoneally injected with a behaviorally-subthreshold dose of clozapine (Caymen Chemical, 0.1mg/kg) to activate the hM4Di receptor at least 10 minutes before the foraging session. The robotic predator was positioned behind the 70cm mark where the food pellet was located. The trial ended after the mouse consumed the pellet and returned to the nesting zone or 5 minutes after the trial began with no successful consumption of the food pellet. Following 24hrs, mice completed two consecutive post-encounter foraging sessions without the presence of the robogator. These post-encounter sessions were identical in structure to baseline foraging. No clozapine was administered in the foraging sessions post-robotic predator exposure.

#### Return to Foraging Baseline

After the post-encounter sessions, mice were allowed to explore the foraging zone and consume a food pellet located at discrete, incremental distances in a similar manner to the initial foraging training. This re-training to baseline foraging was conducted over 5 sessions.

#### Foraging circa- and post-<u>Live Predator Exposure</u>

After a return to baseline foraging, mice were exposed to a live predator in a single session in the last trial, in which the food pellet was located 70cm away from the nesting zone. Mice were intraperitoneally injected with clozapine (0.1mg/kg) at least 10 minutes before the foraging session. The live predator was positioned behind the 70cm mark with the small circular opening pointed towards the nesting zone. The trial ended after the mouse consumed the pellet and returned to the nesting zone or 5 minutes after the trial began with no successful consumption of the food pellet.

#### Foraging post-Live Predator Exposure

Mice completed two consecutive foraging sessions after the live predator encounter. These sessions were identical in structure to baseline foraging. No clozapine was administered in the foraging sessions post-live predator exposure.

#### Statistics

Statistical analyses were performed in R version 4.0.5 (R Core Team, 2021). Groups of 8 mice were used for behavioral analysis to achieve 80% statistical power with an alpha level of 0.05. When multiple variables existed for comparison, data were fitted to a linear mixed effects model such that treatment (GFP, hM4D(Gi)), sex (females, males), session (1-18), trial (30cm, 50cm, 70cm), predator (robogator, live predator), and predator encounter (pre-encounter, circa-encounter, post-encounter) were the fixed effects. Individual mice were treated as a random effect. Linear mixed effects model analyses were conducted using R packages lme4 [35] and lmerTest [36]. These models were evaluated with F-tests on type III sums of squares on the defined fixed effects and their interactions. Post-hoc significance analysis was conducted through comparisons of estimated marginal means using the R package emmeans [37]. Data were fitted to a linear mixed effects model to account for possible unbalanced data and to lower both Type I and Type II errors [38,39]. For analysis of foraging latency by trial, hazard ratios were calculated from a Cox-proportional hazards regression model using the R package survival [40]. For analysis of locomotor linear path before and after predator threat exposure, the y-coordinate (width of the foraging apparatus) for the center point of all subjects was averaged across 10cm-bins of the x-coordinate (length of the foraging apparatus). The averages were used to fit a linear model, in which treatment (GFP, hM4D(Gi)), trial (30cm, 50cm, 70cm), x-coordinate (1-10 bins), and predator encounter (pre-, post-) were the linear predictors. In order to compare the effect of predator exposure and BNST GABA chemogenetic inactivation on multiple foraging parameters, all foraging parameters (foraging latency, last nest exit, first nest exit, distance traveled, nest time, nest exits) were normalized on a scale of 0 to 1 within foraging parameter and trial (30cm, 50cm, 70cm). We defined overall foraging performance as the polygon area calculated from the shape of all parameters plotted on a polar plot. Normalization (((x−min(x))/(max(x)−min(x))=z_(behavior, trial)_) by foraging parameter in addition to trial was due to statistical analysis revealing significant differences in baseline foraging as a result of trial. The polygon area for each subject was calculated to fit a linear mixed effects model and run post-hoc significance analysis, as previously described. When applicable, outliers were detected by the Grubbs test and removed from analysis. All data are reported as mean ± standard error of the mean (SEM), and all significant comparisons are indicated by an asterisk in the figures.

## Acknowledgements

This research was supported by the Webb-Waring Biomedical Research Award from the Boettcher Foundation (DHR), R01 DA047443 (DHR), F31 MH125569 (DJM), seed funds from the Anschutz Boulder Nexus (ABNexus), a 2020 NARSAD Young Investigator grant from the Brain and Behavior Research Foundation (DHR). The imaging work was performed at the BioFrontiers Institute Advanced Light Microscopy Core (RRID: SCR_018302). Laser scanning confocal microscopy was performed on a Nikon A1R microscope supported by NIST-CU Cooperative Agreement award number 70NANB15H226. The funders had no role in study design, data collection and analysis, decision to publish, or preparation of the manuscript. Biorender was used to make select figure panels.

